# Succinct dynamic variation graphs

**DOI:** 10.1101/2020.04.23.056317

**Authors:** Jordan M. Eizenga, Adam M. Novak, Emily Kobayashi, Flavia Villani, Cecilia Cisar, Simon Heumos, Glenn Hickey, Vincenza Colonna, Benedict Paten, Erik Garrison

**Author notes:** Contributed equally (Jordan M. Eizenga, Adam M. Novak).

## Abstract

**Motivation:** Pangenomics is a growing field within computational genomics. Many pangenomic analyses use bidirected sequence graphs as their core data model. However, implementing and correctly using this data model can be difficult, and the scale of pangenomic data sets can be challenging to work at. These challenges have impeded progress in this field.

**Results:** Here we present a stack of two C++ libraries, `libbdsg` and `libhandlegraph`, which use a simple, field-proven interface, designed to expose elementary features of these graphs while preventing common graph manipulation mistakes. The libraries also provide a Python binding. Using a diverse collection of pangenome graphs, we demonstrate that these tools allow for efficient construction and manipulation of large genome graphs with dense variation. For instance, the speed and memory usage is up to an order of magnitude better than the prior graph implementation in the vg toolkit, which has now transitioned to using `libbdsg`’s implementations.

**Availability:** `libhandlegraph` and `libbdsg` are available under an MIT License from https://github.com/vgteam/libhandlegraph and https://github.com/vgteam/libbdsg.

## 1 Introduction

With falling sequencing costs, the genomics community has sequenced increasingly many individuals within certain species. For example, hundreds of thousands of deeply-sequenced human genomes are now available. The novel challenges of analyzing data sets of this scale have led to the development of **computational pangenomics**, which focuses on analyzing populations of genomes rather than individuals (Computational pan-genomics consortium, 2016).

Much of the research in computational pangenomics has coalesced around graph-based approaches for representing populations of genomes (Paten *et al*., 2017). Unlike conventional string-based representations, graph data structures provide a coherent modeling language to represent different types of genomic variation like substitutions, insertions, deletions, and more complex genomic events. They also compactly represent many-way relationships between related genomes, such as whole genome alignments (Kehr *et al*., 2014).

Graph-based data structures also present new computational challenges. In addition to sequence, genome graphs must represent topology. Given the size of many genomes of interest, this can be quite demanding on computer memory. Furthermore, there is significant impetus to make the graph data structures computationally efficient, since they are frequently the core data structure in pangenomics applications.

Genome graphs that include small variants and describe a large population of eukaryotic genomes can contain hundreds of millions of nodes and edges. Using naïve data structures to identify and provide random access to elements of these graphs has very high memory costs. However, the total information content is only incrementally more than in the total sequence set of the pangenome. This suggests that significant memory savings should be possible.

Early versions variation graph toolkit (vg) (Garrison *et al.*, 2018) have provided a cautionary tale of such a naïve implementation. vg used fullwidth machine words as identifiers for graph elements, stored the elements and graph topology in a set of hash tables, and linked identifiers to elements with raw pointers. Loading the 1000 Genomes Project’s variant set into the vg toolkit used to consume more than 300 GB of memory, which is ~30 times as large as the serialized representation (Garrison, 2019).

Although vg provided a memory-efficient succinct representation of the graph (xg) that could be used during read mapping and variant calling, the succinct representation did not allow for dynamic updates to the graph. As a result, graph-modifying steps in vg pangenomic analysis pipelines had to break large graphs into smaller pieces, often connected components that correspond to chromosomes. Unfortunately, this strategy is not tenable for all graphs. For instance, many whole genome alignment and assembly graphs consist of a single giant component that cannot be partitioned easily.

To overcome this limitation, we have developed three new graph genome data structures that are both dynamic, in that they allow efficient updates and edits, *and* succinct, in that they require memory on the order of the graph’s information content. Here, we compare the performance of these data structures to those originally in vg as well as xg using a diverse collection of genome graphs obtained during our work in graphical pangenomics.

In addition to demonstrating the possibility of working with large, complex graphs in small amounts of memory, these implementations expose a common API based on the HandleGraph model described below. This model provides a consistent, reliable interface to genome graphs based on their fundamental elements. The vg toolkit has been refactored to use this API as its default means of serializing and manipulating graphs since version 1.22.0.

We package these implementations behind equivalent C++ and Python APIs in `libbdsg`. This software library will reduce the need for individual research groups to continually reimplement these core data structures. These dynamic HandleGraph libraries will ease the development of algorithms that work on large, complex pangenome graphs by making it easy to store them in reasonable amounts of working memory and manipulate them in reasonable amounts of time.

## 2 Implementation

### 2.1 Data model

Our libraries adopt node-labeled bidirected graphs as a formalism for sequence graphs. In a bidirected graph, nodes are considered to have left and right “sides”, and edges connect two sides rather than two nodes. In bidirected sequence graphs, a node’s sides correspond to the 5’ and 3’ ends of its DNA sequence.

Paths through a bidirected graph must leave a node out of the side opposite the side through which they enter it. We interpret paths that traverse a node from the 3’ side to the 5’ as using the node’s reverse complement strand, which provides a natural means to encode DNA strandedness. Some paths correspond to sequences of interest, such as reference genomes or annotations of the reference. Because paths like these are so frequently important in practice, our formalism also includes paths as a first class object, embedded in the graph.

### 2.2 The HandleGraph interface

The `libhandlegraph` library describes a generic interface that exposes basic operations on our sequence graph data model. The interface uses “handles” (opaque references modeled after the concept of a file handle) in order to remain agnostic about the backing implementation of the graph.

The HandleGraph model focuses on five fundamental entities in bidirected sequence graphs (Figure 1):

- *Nodes* identify pairs of complementary DNA strands and have unique numerical identifiers (IDs).
- *Strands* identify one strand of a node’s DNA sequence.
- *Edges* link pairs of strands, in order.
- *Paths* represent sequences as walks through the graph.
- *Steps* describe paths’ visits to nodes’ strands.

**Fig. 1.**
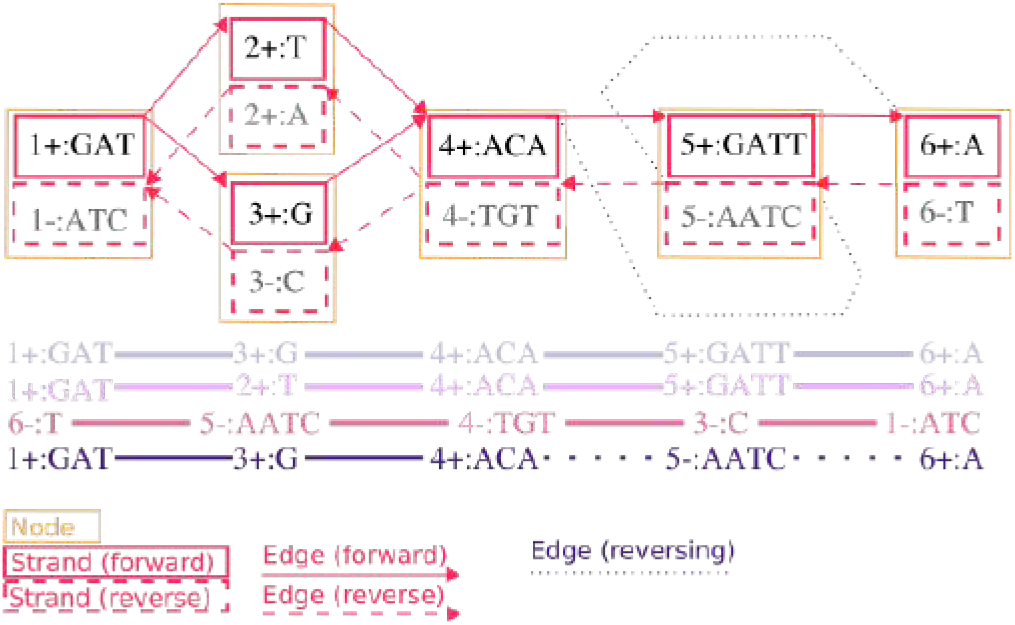
Entities in the bidirected sequence graph. Top: a variation graph showing nodes (yellow rectangles), each of which contain a forward and reverse strand (red solid and dashed rectangles, respectively). Strands show the node identifier, the direction (+ or —), and the sequence of the strand. Note that reverse strands show the reverse complement sequence of the forward strand. All edges are shown as connections between nodes, with forward-to-forward edges denoted by solid lines, while reverse-to-reverse edges denoted by dotted lines. Two edges that invert from forward to reverse and reverse to forward are shown with dotted lines. Edges run from the strand at their beginning to their end, as indicated by the arrowhead. Bottom: an illustration of four paths. (Each has an implicit handle, and a name, which are omitted for brevity.) Each path is shown in its natural direction as a series of connected steps that refer to strands in the graph. The first two paths differ by a SNP, with one passing through `2+:T`, and the other through `3 +:G`. The third path is the reverse complement of the first. The fourth is the same as the first, but contains an inversion, passing through `5-:AATC` rather than `5+:GATT`.

The defining feature of the model is that none of these entities are accessed directly. Instead, only *handles*, or references, to strands, edges, paths, and steps are available. The `libhandlegraph` interface requires sequence graph implementations to be able to provide or consume these handles for all queries. For instance, we might obtain a handle to a strand from an implementation by querying a node’s ID and providing an orientation, while another function provided in the implementation would map the handle back to a given node ID. Alternatively, we can obtain a handle to a path from its name (e.g. “chr22”), and then iterate over handles to the path’s steps to follow its course through the graph. Or, starting from a given handle to a strand, we can follow its outgoing or incoming edges to explore the topology of the graph.

The actual contents of a handle are unspecified, which gives significant flexibility to the implementation. One benefit of this design is that any algorithm designed for one HandleGraph implementation can be applied to all other implementations. Another is that, since the user works only through handles that they cannot forge or modify, their ability to make mistakes can be restricted. For example, the interface can enforce the constraints on paths through bidirected graphs during edge traversal. Furthermore, the interface is able to be memory-safe by eliminating raw pointers and other direct access to graph elements.

### 2.3 Graph implementations

We consider five implementations of the HandleGraph model. To ground our experimental results, here we we provide a high-level overview of each implementation. Two implementations, vg and xg, have been described previously (Garrison *et al*., 2018; Garrison, 2019). The others are combined in the `libbdsg` library (https://github.com/vgteam/libbdsg), which provides three concrete implementations: HashGraph, odgi, and PackedGraph. Each implementation represents a different tradeoff in terms of speed, memory use, and capabilities. All of the implementations except xg are dynamic. Since all of these implementations use the same interface, the `libhandlegraph` header files serve as the most effective developer documentation for their functionality.

#### 2.3.1 vg

We have extended the data model in vg, previously described in (Garrison *et al*., 2018), to match the HandleGraph API. The backing data structures used by the model remain the same. The graph entities are stored as objects in a backing vector, and referred to internally by hash tables that map between node identifiers and pointers into this vector. Edges are indexed in a hash table mapping pairs of handles to edge objects. Paths are stored in a set of linked lists, with a hash table mapping between nodes and path steps. This arrangement was tenable for the early development of algorithms working on variation graphs. Its inefficiency, caused by unnecessary overheads and data duplication, has generated significant difficulty for groups working with vg. The other HandleGraph implementations respond to the limitations of this approach. In version 1.22.0, vg was updated to use HashGraph (below) as the default format, though it remains compatible with all implementations described in this paper via the HandleGraph API.

#### 2.3.2 xg

xg was initially developed in response to the memory and runtime costs of vg, which prevent its application to large graphs. It additionally provides positional indexes over paths that are required for read mapping and variant calling, and is the graph data model used in most established bioinformatic operations on variation graphs (Garrison *et al*., 2018; Hickey *et al*., 2020). Unlike other HandleGraph implementations, xg is a static graph index. This permits a more powerful set of efficient queries against the graph, especially for paths. The encoding is designed to balance speed and low memory usage. The topology of the graph is encoded in a single vector of bit-compressed integers, which promotes cache-efficiency. Rank and select operations on succinct bit vectors are used to provide random access over the variable-length records, which each encode a node’s sequence, ID, and edges. Embedded paths are encoded in variable-length integer vectors with Elias gamma encoding. Rank and select operations on succinct bit vectors also provide queries by base-pair position along paths.

#### 2.3.2 HashGraph

HashGraph has speed as its primary goal. It represents the graph as a collection of node objects in a high performance hash table, while embedded paths are implemented as doubly-linked lists. Edges are recorded in vectors attached to each node that they connect. This adjacency list encoding is appropriate for genome graphs, which are typically very sparse. Each node object maintains a vector of pointers to the path steps that traverse it. Most of HashGraph’s component data structures are uncompressed STL objects which can be used efficiently in their native in-memory arrangement. HashGraph trades memory for time, and thus is most appropriate for small graphs (from small genomes or small regions of larger genomes) or for high-memory compute environments.

#### 2.3.4 odgi

odgi (Optimized Dynamic Graph Implementation) is based on a node-centric encoding of the graph that is designed to improve cache coherency when traversing or modifying the graph. This encoding is split between graph topology and paths, which is important for achieving a good balance of runtime performance and memory usage on real-world graphs with large path sets. Each node’s sequence and edges are encoded in a byte array using a variable-length integer encoding scheme. Edges are described in terms of a relative offset between the rank of this node in the sorted array of nodes of the graph and the node to which the edge arrives. The set of steps traversing the node is recorded in a second dynamic integer vector, compressed so that only the largest integer entry is stored at full bit-width (Prezza, 2017). Each step contains a path identifier, relative ranks of the previous and next nodes on the path, and the ranks of the previous and next steps among the path steps recorded at their nodes. This path encoding scheme is similar to that used in the dynamic GBWT (Sirén *et al*., 2020), but differs in that the paths are not prefix-sorted. If each path step tends to move only a short distance in the sorted nodes of the graph (e.g. from *n*_5_ → *n*_7_), then the maximum bit-width of the path vector will be low, resulting in good compression.

#### 2.3.5 PackedGraph

PackedGraph is designed to have a very low memory footprint. Most of its component data structures are—conceptually speaking—linked lists. However, they are implemented using vectors of bit-compressed integers, where pointers are produced by treating some of the integer entries as indexes into the vector that contains them. The bit-width can be determined dynamically. Doing so does not affect the amortized asymptotic run time of graph operations in the typical case that the value of *i*-th entry in the vector is *O*(*i*). The vector uses a windowed bit compression scheme in which only one value within a window is maintained at its full bit-width. The remaining entries are represented as differences from this value. In the typical case where adjacent entries in the vector are highly correlated, this helps keep the bit-width low and the compression high.

### 2.4 Python binding

We have implemented a Python binding to the graph implementations in `libbdsg` using Pybind11 (Jakob *et al*., 2017). This allows the data structures to be used in Python applications, significantly lowering the barrier-to-entry for pangenomic application developers. This functionality is documented at https://bdsg.readthedocs.io, including a tutorial. This documentation also serves as useful introduction to the HandleGraph API.

### 2.5 Code availability

Both `libhandlegraph` and `libbdsg` are open source under an MIT License. They are available on GitHub at https://github.com/vgteam/libhandlegraph and https://github.com/vgteam/libbdsg. Extensive documentation of these libraries and their respective graph implementations is available at https://pangenome.github.io/.

## 3 Evaluation

### 3.1 Human genome with structural variants

We measured the core operation performance of the four graph implementations and the graph class from the popular vg software (as implemented prior to version 1.22.0). In particular, we measured 1) memory usage to construct a graph, 2) time to construct a graph, 3) memory usage to load an already-constructed graph, and 4) time to access nodes, edges, and steps of a path. The presented results are from a graph describing the structural variants of the Human Genome Structural Variation Consortium (Chaisson *et al*., 2019). The results generally match our expectations based on the implementations’ design goals (Figure 2).

**Fig. 2.**
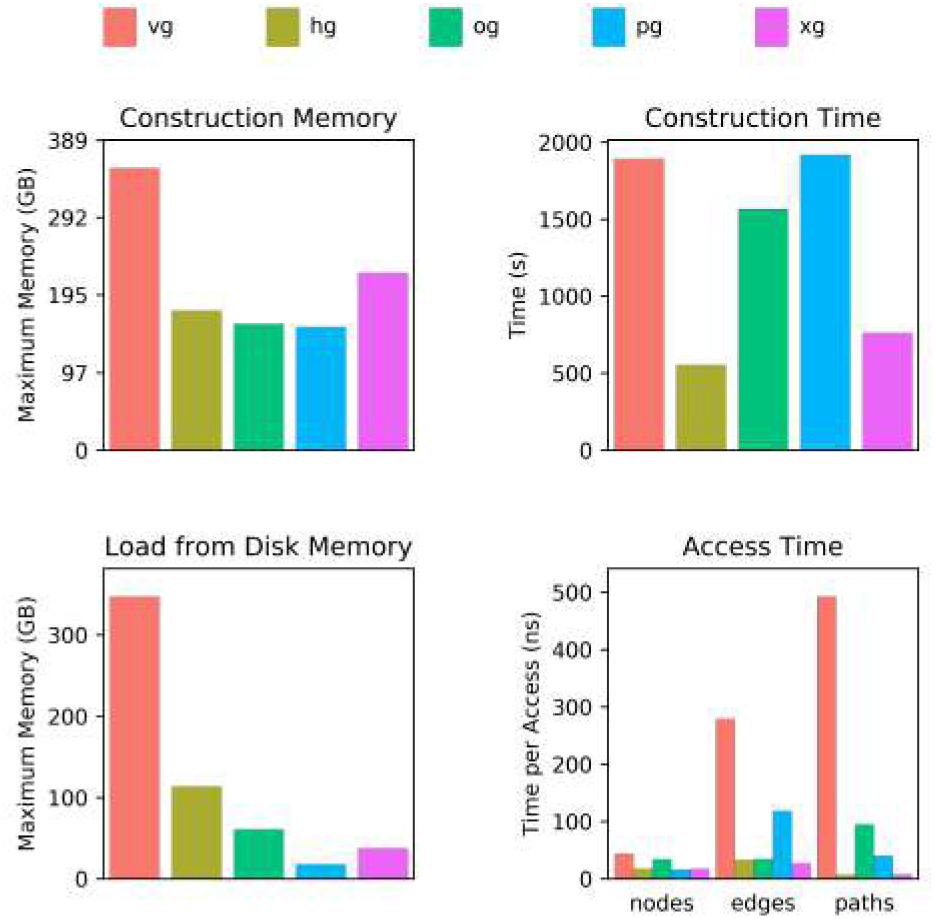
Performance on a graph of structural variants from the HGSVC. Abbreviations used here and in subsequent figures and tables: vg = vg, hg = HashGraph, og = odgi, pg = PackedGraph, xg = xg. All four new graph implementations compare favorably to vg. PackedGraph tends to be the most memory efficient, HashGraph tends to be the fastest, and odgi is balanced in between. xg provides good performance on both memory usage and speed, but it is static.

### 3.2 Genome graph collection

To compare the methods’ performances across a wide variety of different graphs, we applied each to a collection of 2299 graphs collected during our research on graphical pangenomics. For each graph and graph implementation, we measured the same metrics described in the previous section as well as various graph properties including size, edge count, cyclicity, and path depth. We summarize these results in Figures 3 and 4.

**Fig. 3.**
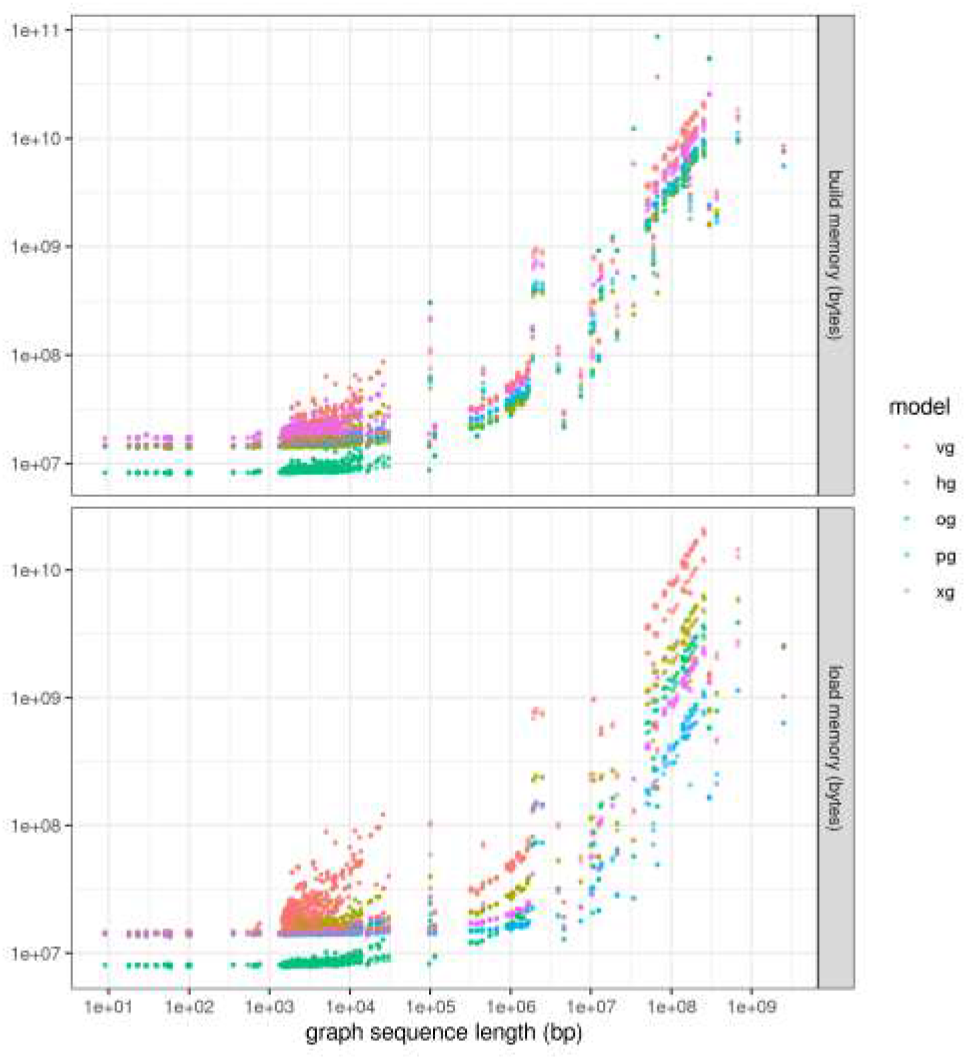
Memory requirements for model construction and loading. Memory costs versus graph sequence size for the graph collection, colored by HandleGraph model. The memory requirements for graph construction tend to be higher than those for loading the graph model. All methods show fixed overheads of several megabytes, seen in the flat tail to the left of both plots. Outside of this region, all methods show roughly linear scaling in both build and load costs per input base pair. The relative differences in memory costs appear to be stable between different methods across many orders of magnitude in graph size. Notably, vg has approximately the same build and load costs, as the data structures used are the same. The other methods tend to use auxiliary data structures at build time, and so require more memory to build the data structure than to load it.

**Fig. 4.**
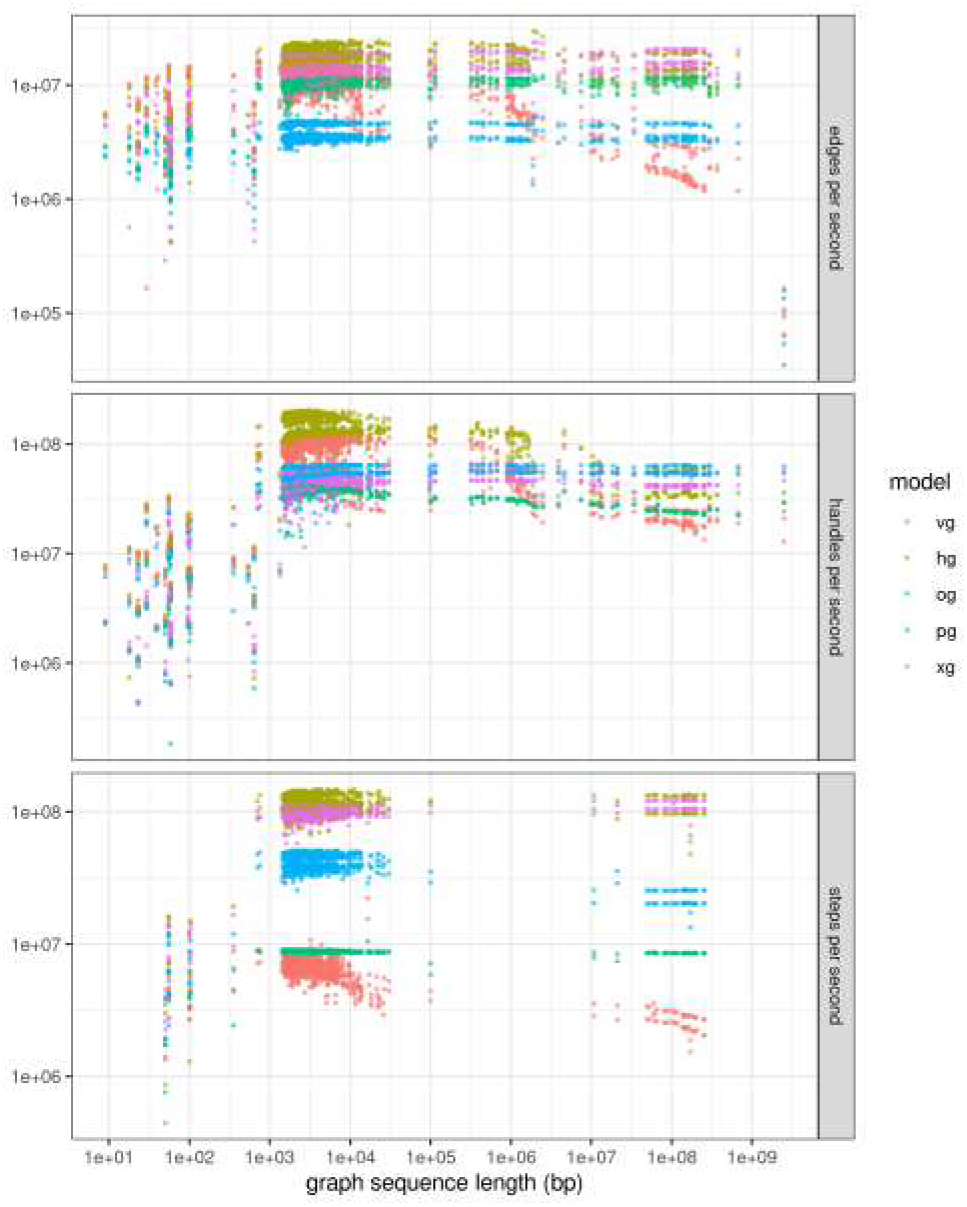
Graph element enumeration performance. Iteration performance for edges, nodes, and path steps for the full graph collection, shown in terms of elements per second. HashGraph provides the highest performance for all element iteration types on smaller graphs, but this performance falls of with larger graphs, presumably due to scaling properties of the backing hash tables. The same pattern can be seen for vg, although the overall performance is worse. Although it has the worst edge iteration performance, PackedGraph provides good performance on node and path step iteration. The relative path encoding in odgi yields poor performance on path iteration, and node decoding overheads appear to reduce its node iteration performance, but it has good graph topology traversal performance, perhaps due to cache coherency of the edge encoding. xg provides excellent iteration performance in all cases.

For graph construction and loading, we observe similar trends as for the HGSVC graph. vg’s performance in terms of memory usage is very poor, both during construction and load. For construction and load, all models exhibit largely linear scaling characteristics, outside of very small graphs where static memory overheads dominate. PackedGraph yields the best memory performance for larger graphs (which are mostly the chromosomes of the 1000 Genomes Project graph), while for the mediumsized graphs in the collection (~1 Mbp), odgi requires less memory.

For graph queries and iteration, the relative performance of the models is largely maintained across the entire range of graph sizes. However, we observe that the hash-based models (vg and HashGraph) have very good performance for smaller graphs (in handle and edge enumeration) but decrease in throughput as the graph size increases. Smaller, less dramatic decreases in performance can be seen for the other implementations. For path enumeration, the highest-performing methods are xg and HashGraph at approximately 10 times faster than odgi, whose relative path storage is costly to traverse.

### 3.3 1000 Genome Project chromosome graphs

Variation graphs built from the 1000 Genomes Project (1000GP) variant catalog and the human reference genome have fairly homogenous and regular features. In addition, they have connected components of very different sizes, each corresponding to a chromosome. This provides a natural, fairly controlled means to explore the scaling behavior of our data structures. Moreover, graphs of this form are seeing increasing use in variant-aware resequencing analyses (Crysnanto and Pausch, 2019). Thus, the performance of data structures on these graphs is of general interest.

We first evaluated the scaling performance of the various HandleGraph implementations relative to node count for each of the nuclear chromosomes in the 1000GP (Figure 5). We find that for all methods, load memory scales almost perfectly with node count, with an average *R*^2^ = 0.998. Due to differences in variant density among the chromosomes, the average correlation relative to sequence length is lower (*R*^2^ = 0.986).

**Fig. 5.**
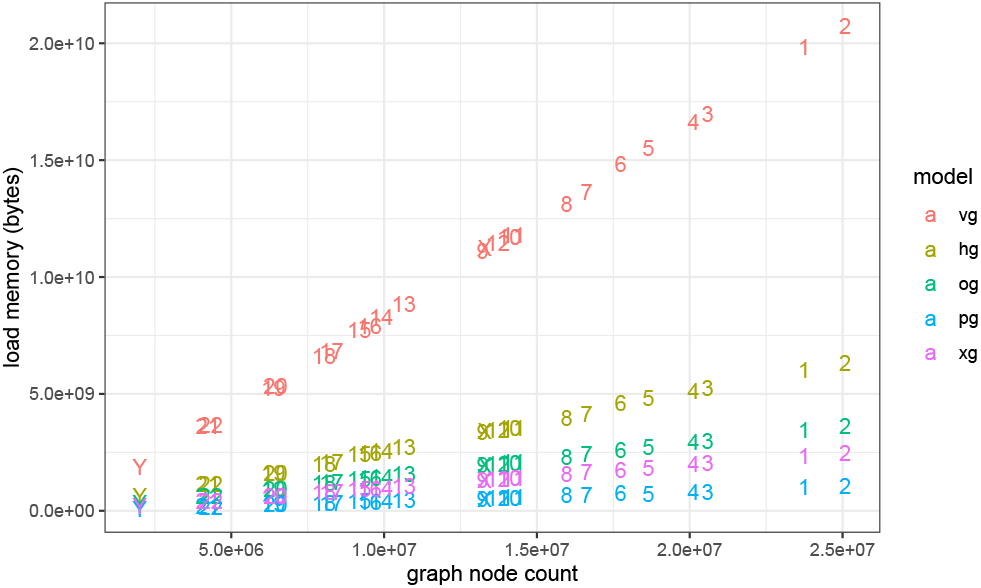
Load memory versus node count for chromosome graphs built from 1000 Genomes Project variants and GRCh37. For each method, memory requirements are more strongly correlated with the number of nodes in the graph (*R*^2^ = 0.998) than with the graph sequence length (*R*^2^ = 0.986). Although the memory requirements are dominated by graph sequence size, node count will increase with variant density. Methods generally incur an overhead for each node that is larger than the sequence length. Linear scales clarify that the absolute difference in performance between vg and the other methods is significant.

In Table 1, we report the average memory performance of the methods relative to graph sequence length, and also the iteration performance in terms of elements per second. We find that the best-performing method in terms of memory usage is PackedGraph, which consumes around 1/20th the memory of vg per base-pair of graph in the 1000GP set. However, it provides significantly better iteration performance for nodes, edges, and path steps. HashGraph and xg have similar iteration performance, but xg, by virtue of its use of compressed, static data structures, requires less than half as much memory. odgi optimized for efficient dynamic operations on graphs with higher path coverage, and in general is not as performant as other methods on this set.

**Table 1.**
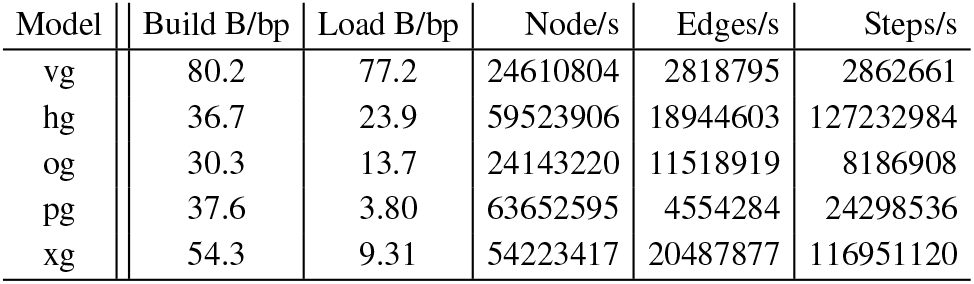
Performance on 1000 Genomes Project chromosome graphs. Average build memory, load memory, and iteration times for graph elements for the chromosome-level graphs built from all the variants in the 1000 Genomes Project and the GRCh37 reference genome against which the variant set was originally reported. vg requires ~ 20 times as much memory to load the graphs as PackedGraph, while even the most costly HandleGraph model (HashGraph) requires ~ 1/3 as much memory. In these graphs, odgi provides the lowest performance for handle iteration. However, in all other metrics, vg performs significantly worse than the other models.

| Model | Build B/bp | Load B/bp | Node/s | Edges/s | Steps/s |
| --- | --- | --- | --- | --- | --- |
| vg | 80.2 | 77.2 | 24610804 | 2818795 | 2862661 |
| hg | 36.7 | 23.9 | 59523906 | 18944603 | 127232984 |
| og | 30.3 | 13.7 | 24143220 | 11518919 | 8186908 |
| pg | 37.6 | 3.80 | 63652595 | 4554284 | 24298536 |
| xg | 54.3 | 9.31 | 54223417 | 20487877 | 116951120 |

## 4 Discussion

We have presented a set of simple formalisms, the HandleGraph abstraction, which provides a coherent interface to address and manipulate the components of a genome variation graph. To explore the utility of this model, we implemented data structures to encode variation graphs and matched them to this interface. This allowed us to directly compare these HandleGraph implementations on a diverse set of genome graphs obtained during our research. These experiments reveal that genome graphs need not pay the computational expense of the early versions of vg. The best-performing models require an order of magnitude less memory than vg while providing higher performance for basic graph access operation and element iteration. For these reasons, vg has transitioned to using these newer graph implementations.

The efficiency of these methods and their encapsulation within a coherent programming interface will support their reuse within a diverse set of application domains. Variation graphs have deep similarity with graphs used in assembly; these libraries could be used as the basis for assembly methods. They could also be used for genotyping and haplotype inference methods based on graphs (Garg *et al*., 2018).

Ongoing work is establishing large numbers of highly-contiguous whole genome assemblies for humans (https://humanpangenome.org/). Improvements in sequencing technology are likely to make such surveys routine. It is natural to consider a pangenome reference system, based on the whole genome alignments of such assemblies, as the output of these pangenome projects. Recent results demonstrate that many basic bioinformatic problems can be generalized to operate on such structures. Should these pangenome representations become common or standard, then variation graph data structures like those we have presented here will form the basis for a wide range of pangenomic methods.

